# The molecular basis of *Acinetobacter baumannii* cadmium toxicity and resistance

**DOI:** 10.1101/2020.10.20.348086

**Authors:** Saleh F. Alquethamy, Felise G. Adams, Ram Maharjan, Natasha N. Delgado, Maoge Zang, James C. Paton, Karl A. Hassan, Ian T. Paulsen, Christopher A. McDevitt, Amy K. Cain, Bart A. Eijkelkamp

## Abstract

*Acinetobacter* species are ubiquitous Gram-negative bacteria that can be found in water, soil and as commensals of the human skin. The successful inhabitation of *Acinetobacter* species in diverse environments is primarily attributable to the expression of an arsenal of stress resistance determinants, which includes an extensive repertoire of metal ion efflux systems. Although metal ion homeostasis in the hospital pathogen *Acinetobacter baumannii* is known to contribute to pathogenesis, insights into its metal ion transporters for environmental persistence are lacking. Here, we studied the impact of cadmium stress on *A. baumannii*. Our functional genomics and independent mutant analyses revealed a primary role for CzcE, a member of the cation diffusion facilitator (CDF) superfamily, in resisting cadmium stress. Further, we show that the CzcCBA heavy metal efflux system also contributes to cadmium efflux. Analysis of the *A. baumannii* metallome under cadmium stress showed zinc depletion and copper enrichment, which are likely to influence cellular fitness. Overall, this work expands our understanding of the role of membrane transporters in *A. baumannii* metal ion homeostasis.

**IMPORTANCE:** Cadmium toxicity is a widespread problem, yet the interaction of this heavy metal with biological systems is poorly understood. Some microbes have evolved traits to proactively counteract cadmium toxicity, which includes *Acinetobacter baumannii*. Here we show that *A. baumannii* utilises a dedicated cadmium efflux protein in concert with a system that is primarily attuned to zinc efflux, to efficiently overcome cadmium stress. The molecular characterization of *A. baumannii* under cadmium stress revealed how active cadmium efflux plays a key role in preventing the dysregulation of bacterial metal ion homeostasis, which appeared to be the primary means by which cadmium exerts toxicity upon the bacterium.

## INTRODUCTION

The heavy metal cadmium is enriched in industrial settings and mining sites through anthropogenic activity. Although various metalloproteins retain their function when bound to cadmium (1), cadmium is primarily associated with toxicity in biological systems (2). Despite being redox inactive, cadmium stress is commonly connected to the induction of oxidative stress. Recent advances in the field have revealed cadmium-metalloprotein interactions as a primary feature of cadmium toxicity (3, 4). In addition to dysregulation of central carbon metabolism and lipid homeostasis, cadmium stress in *Streptococcus pneumoniae* causes manganese and zinc starvation via interaction with non-cognate metalloproteins, these being a manganese-recruiting protein and a zinc-responsive transcriptional regulator (4). Manganese is the primary co-factor for superoxide dismutase in *S. pneumoniae*, hence, cadmium-induced manganese depletion leads to oxidative stress susceptibility (4, 5). In *Salmonella enterica*, excess cadmium was found to affect zinc and manganese accumulation, but not iron (3). These studies exemplify that dysregulation of metal ion homeostasis and protein mismetallation by cadmium is central to its toxicity in cells.

*Acinetobacter baumannii* is rapidly emerging as one of the world’s most antimicrobial resistant bacterial pathogens (6–8). Its highly effective intrinsic and acquired stress resistance features have allowed this human pathogen to survive in the hospital environment, and thrive when colonising immunocompromised individuals (9, 10). Various members of the *Acinetobacter* genus, including some strains of *A. baumannii*, have a strong environmental presence. This has resulted in the *A. baumannii* genome harbouring diverse metabolic and stress resistance features that are typically not linked to pathogenicity (11, 12). Its environmental roots are also reflected in the extensive arsenal of putative heavy metal export mechanisms that may enable persistence under diverse metal ion stresses (13, 14). *A. baumannii* metal ion biology is a rapidly expanding field of research. In particular its diverse array of metal ion acquisition systems have been studied intently (15–21). In contrast, our understanding of heavy metal resistance is limited, with only the roles of zinc and copper toxicity being studied in recent years (13, 14, 22–24). Metal ion toxicity analyses in *A. baumannii* have revealed widespread implications on cellular physiology, including dysregulation of metal ion homeostasis, oxidative stress susceptibility and alterations in membrane biology. Similar to the impact of cadmium on the dysregulation of *S. pneumoniae* and *S. enterica* metal ion homeostasis, zinc stress in *A. baumannii* has been shown to result in copper depletion (13).

The *A. baumannii* genome harbours a broad arsenal of both highly-conserved and recently-acquired efflux systems that have putative functions in metal resistance (13). The P-type ATPase CopA has been shown to be the primary mediator of cytoplasmic copper efflux in *A. baumannii* and plays a role in its virulence potential (23). The complexity of copper homeostasis in the periplasm has also been studied (24), but strain-to-strain variation complicates defining their significance across the species (13, 23). Zinc resistance is mediated by an efficient efflux pathway which consists of the cation diffusion family (CDF) member CzcD, a periplasmic zinc chaperone named CzcI and the heavy metal efflux (HME) system CzcABC (22).

Bacterial cadmium efflux can be mediated by HME, CDF or P-type ATPase family members. Examples include CadA, a P-type ATPase from *Streptococcus thermophilus* (25), the CDF member YiiP from *Escherichia coli* (26, 27) and the CznABC HME system from *Helicobacter pylori* (28). These systems may display overlapping substrate profiles within a bacterial species, which requires experimental verification. Although a recent study has provided insights into the role of the C-terminal domain in CDF metal specificity (29), overall, the specificity for transition metals varies greatly between members within the CDF, HME and P-type ATPase families and is difficult to predict based on protein sequence analyses.

In this study, we define the genome-wide cadmium stress adaptation mechanisms in *A. baumannii*. This work identified both CDF and HME systems that play key roles in cadmium export in *A. baumannii* which was confirmed by single-gene mutant analyses. We also show that the bacterium encodes a highly efficient cadmium sensing regulator for the transcriptional activation of the primary cadmium resistance mechanism. Through the global metallomic analyses under cadmium stress, this study provides novel insights into cadmium toxicity.

## METHODS

### Bacterial strains, chemicals, media and growth conditions

*Acinetobacter baumannii* AB5075_UW mutant derivatives were purchased from the Manoil Laboratory (30) and T26 transposon insertions confirmed by PCR. The *A. baumannii* ATCC 17978 *czcA* and *czcE* mutants were generated in our laboratory for a previous study (22) (See **Tables S1** and **S2** for list of strains and oligonucleotides used in the study, respectively). All chemicals were purchased from Sigma Aldrich (Australia) unless otherwise indicated.

*A. baumannii* strains were routinely grown in Luria Bertani broth (LB), containing 1% tryptone (BD Bacto), 0.5% yeast extract (BD Bacto) and 0.5% sodium chloride. For overnight culturing, a single colony from LB agar (1.5%) was used to inoculate 4 mL of LB medium. Overnight cultures were diluted to an optical density at 600 nm (OD_600_) of 0.01 in 200 μL LB for growth assays or 20 mL LB for all other analyses. For growth assays, cultures in LB media were incubated at 37°C with shaking in a FLUOstar Omega Spectrophotometer (BMG Labtech), with the OD_600_ values presented. The 20 mL cultures used for all other analyses were incubated at 37°C in an Innova 40R shaking incubator (Eppendorf) at 230 rpm until they reached an OD_600_ of 0.7.

### Construction of transposon mutant library

*A. baumannii* ATCC 17978 was used to generate a dense transposon library. The library was constructed using the protocol as previously described (31). Briefly, transposomes were prepared using an EZ-Tn*5* transposase (Epicentre Biotechnology) and a custom transposon carrying a kanamycin resistance cassette amplified from the pUT_Km plasmid (32) using the primer sets listed in **Table S2**. The transposomes (0.25 μL) were electroporated into 60 μL of freshly prepared electrocompetent *A. baumannii* cells using a Bio-Rad GenePulser II (1.8 kV, 25 μF, and 200 Ω) in a 1 mm electrode gap cuvette (Bio-Rad). Cells were recovered by resuspension in 1 mL of SOC medium and incubated at 37°C with shaking (200 rpm) for 2 h. Tn*5* insertion recombinants were selected on Mueller Hinton (MH) agar (BD Bacto) supplemented with 7 μg.mL^-1^ kanamycin. Approximately 12 to 16 transformations were performed and pooled for each batch, with 10,000 to 50,000 resulting transformants. Approximately 250,000 mutants were collected and pooled from a total of 10 batches and snap-frozen and stored in 20% glycerol at −80°C.

### Time kill assay for the selection of cadmium concentration for mutant library treatment

Approximately 10^9^ CFU from an overnight ATCC 17978 culture was sub-cultured into fresh 10 mL MH broth spiked with different CdCl_2_ concentrations (0, 3, 60, 80 160, and 320 μM) and incubated at 37°C with shaking. At 0, 1, 2, 4, 5 and 24 h time points, 100 μL samples were taken, 10-fold serially diluted in sterile phosphate-buffered saline (PBS) and 10 μL of each dilution spotted on MH agar. Colonies were enumerated to determine the surviving cells after overnight incubation at 37°C.

### Transposon-directed insertion site sequencing (TraDIS) of mutant library and data analysis

Approximately 10^9^ viable Tn*5* mutant cells were inoculated into 10 mL of MH broth (BD Bacto) and grown at 37°C for 8 h with shaking (200 rpm). A total of 500 μL containing approximately 10^9^ cells were seeded into 10 mL fresh MH broth with or without 60 μM CdCl_2_ in duplicate and grown for 16 h at 37°C with shaking (200 rpm). Genomic DNA was then extracted from approximately 10^10^ cells using the DNeasy UltraClean Microbial Kit (Qiagen) according to the manufacture’s protocol. Sequencing and analysis of the transposon mutant library were performed using the transposon-directed insertion site sequencing (TraDIS) protocol as described previously (33). The primer sets used for PCR amplification of TraDIS fragments (Pf5_PCR) and sequencing (Pf5_Seq) are listed in **Table S2**. Samples were sequenced on a HiSeq2500 Illumina sequencing platform, generating approximately 2 million 50 bp single-end reads per sample as previously described (34). TraDIS sequence reads were deposited in the European Nucleotide Archive under accession number ERP118051 and analysed using the BioTraDIS pipeline with default parameters (as described in (33)).

### Cellular metal ion content analysis

Untreated and metal-stressed bacteria (LB supplemented with 1 μM CdCl_2_ or 15 μM CdCl_2_) were harvested at mid log-phase (OD_600_ 0.7) and washed, by resuspension and centrifugation at 7,000 × *g* for 8 mins, first with PBS containing 5 mM ethylenediaminetetraacetic acid (EDTA), and then with PBS as described previously (22, 23, 35, 36). Bacterial pellets were desiccated at 95°C overnight, followed by determination of dry cell weight, and resuspension of pellets in 35% HNO_3_ and boiling at 95°C for 1 h. Samples were diluted to a final concentration of 3.5% HNO_3_ and analysed by inductively coupled plasma-mass spectrometry (ICP-MS) on an Agilent Solution 8900 QQQ ICP-MS (Adelaide Microscopy, University of Adelaide).

### qRT-PCR

For RNA extraction and qRT-PCR analysis, untreated or metal-stressed bacteria (LB supplemented with 30 μM CdCl_2_) were harvested when they reached mid-log phase (OD_600_ 0.7) and cells lysed in QiaZol (Qiagen) as described previously (37, 38). RNA was extracted and purified using a PureLink RNA Mini Kit (ThermoFisher Scientific), with on-column DNase I treatment, according to the manufacturer’s instructions. qRT-PCR was performed using the SuperScript III One-Step RT-PCR kit (ThermoFisher Scientific) on a QuantStudio 7 Flex System (ThermoFisher Scientific). Transcription levels were corrected to those obtained for *GAPDH* prior to normalization to the transcription levels observed for untreated *A. baumannii* cultures. Oligonucleotide sequences are listed in (**Table S2**).

### Bacterial lipid extraction and gas chromatography/mass spectrometry analysis

Overnight cultures of *A. baumannii* AB5075_UW were diluted to an OD_600_ of 0.05 in fresh LB media (20 ml) and grown to mid-log phase (OD_600_ = 0.7), with treated cultures supplemented with 15 μM CdCl_2_. Cells were harvested by centrifugation at 7,000 × *g* for 10 minutes and washed once with PBS, processed pellets were then resuspended in 50 μl of 1.5% NaCl buffer. For lipid extraction, 1 ml of chloroform:methanol (2:1; v/v) was added to the cell suspension, mixed vigorously for 2 minutes then incubated at RT for 10 minutes. Following the addition of 200 μl 1.5% NaCl, the cell suspension was mixed vigorously for 1 minute and centrifuged at 6,000 × *g* for phase-separation. The lower phase was recovered and concentrated via nitrogen evaporation. All samples were stored at −20°C prior to gas chromatography / mass spectrometry (GC/MS) analysis.

To generate fatty acid methyl esters (FAMEs), concentrated lipid samples were resuspended in 1:1 chloroform and trimethylsulfonium hydroxide (TMSOH) and subsequently analysed using an Agilent 7890A GC system with a 30 m Agilent DB-FastFAME column (Agilent Technologies). Mass spectrometry was completed using a coupled Agilent 5975C MSD system (Agilent Technologies). FAME species were differentiated and determined by comparing to the FAME mix c4-24 standard (Sigma Aldrich). Data analysis was completed using the Agilent MassHunter Qualitative Navigator software (Agilent Technologies).

## RESULTS AND DISCUSSION

### *A. baumannii* employs multiple transport systems to resist cadmium stress

To delineate the molecular basis of cadmium toxicity and the mechanisms that aid in cadmium stress resistance we created a Tn*5*-based random transposon mutant library in the laboratory reference strain, *A. baumannii* ATCC 17978. TraDIS sequencing of the base library in rich media revealed a density of at least 113,000 unique mutants. We subjected 10^9^ mutant cells to a subinhibitory concentration (60 μM) of cadmium for 16 h. This concentration was found to present a minor but significant impact on proliferation of this strain (**Fig. S1**), and thus deemed appropriate for TraDIS analysis. Cadmium exposure resulted in 56 genes displaying significantly differential numbers of insertions (**Table S3**): decreased mutant fitness (decreased transposon insertions) in 14 genes and enhanced fitness (increased transposon insertions) in 42 genes (fold change of ≥ 2; *Q* value, ≤ 0.05).

The TraDIS analyses identified ACX60_13165, which encodes a putative MerR-like transcriptional regulator, as most affected by cadmium treatment (−4.1 fold change) (**Fig. 1**). The regulatory gene is divergently transcribed to the CDF family transport gene, *czcE* (ACX60_13160), which our analyses defined as the gene with the second greatest loss of represented transposon mutants under cadmium stress (−3.8 fold change). The previously characterized HME-RND zinc efflux system, CzcCBA (ACX60_01405-15) and its metal chaperone *czcI* (ACX60_01400) (22), were also shown to be important in conferring cadmium resistance, where mutant abundance decreased between −2.4 and −3.0 fold change. The CDF member, *czcD*, previously revealed to play a role in zinc efflux, was not found to be significantly impacted by cadmium treatment. We identified that transposon mutants of putative DmeF-like cobalt exporting CDF (ACX60_18165) were enriched under cadmium stress (2.1 fold change). This indicates that cobalt may be accumulated at reduced levels under cadmium stress and its undesired export by this DmeF-like transporter is subsequently detrimental to *A. baumannii* fitness.

**Figure 1.**
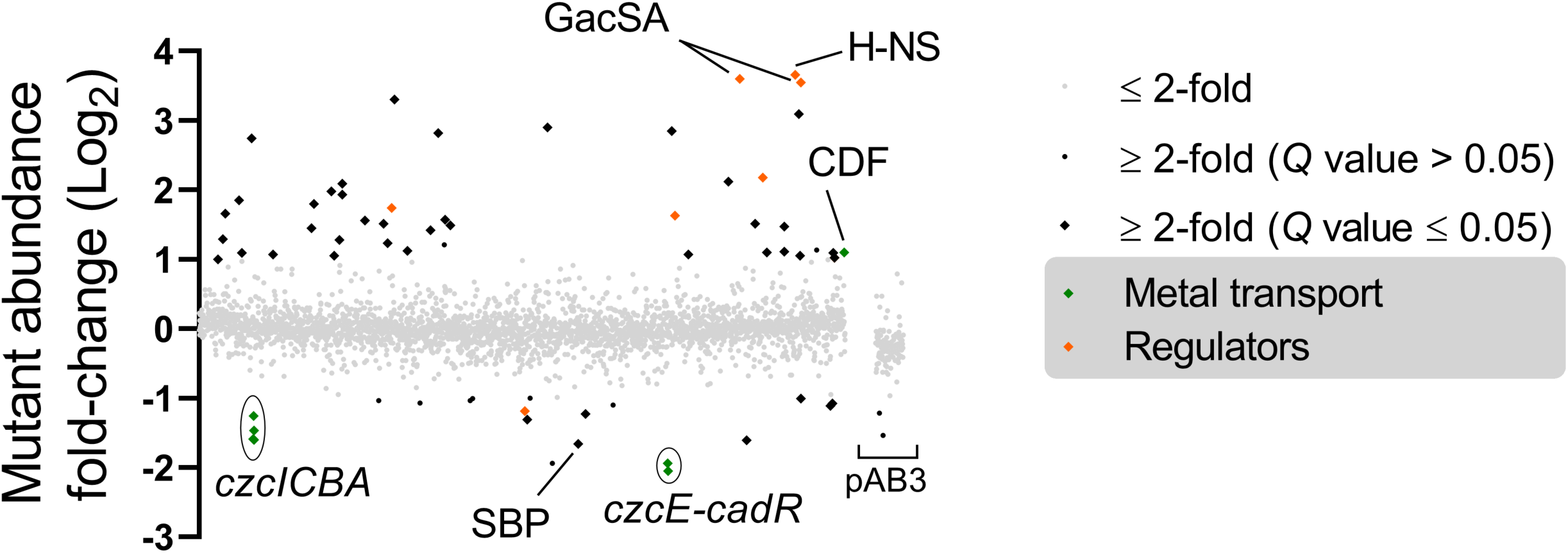
The *A. baumannii* cadmium resistome. The effect of cadmium treatment (60 μM for 16 hours) on the abundance of gene mutations (Tn*5* insertions) mapping to the *A. baumannii* ATCC 17978 chromosome and plasmid pAB3 as determined by TraDIS analysis. The data represent the mean of biological triplicates. The *Q* value was determined by using the method described previously (33).

A solute binding protein (SBP; ACX60_10710) with similarity to methionine and metal ion binding SBPs such as NlpA (COG1464.1), was impacted by cadmium stress (−3.2 fold change). The gene encoding this periplasmic protein is located in an operon that also encodes its putative ATP-binding cassette (ABC) transporter for cargo delivery and subsequent translocation across the inner membrane, but none of these genes were affected by cadmium stress. Hence, the SBP may play a role in periplasmic metal buffering instead of facilitating transport.

The relatively large number of genes in which insertional disruption proved to impart a positive effect included various transcriptional regulators. H-NS (12.6 fold change) is one of *A. baumannii’*s most prominent global transcriptional regulators (39). Although, no role in metal ion stress has been indicated in previous transcriptomic analyses of an ATCC 17978 *hns* mutant, in *Salmonella*, the ferric uptake regulator (FUR) regulates the expression of *hns* and H-NS competes with FUR for some gene targets in *Salmonella* (40, 41). The GacAS regulatory system, of which mutation is beneficial to survival during cadmium stress (11.7 and 12.1 fold change, respectively), plays a key role during infection and in lifestyle changes, including biofilm formation (42). Other regulators include those with putative roles in sensing saturated fatty acids, including ACX60_05275 (3.3 fold change) and ACX60_17890 (2.0 fold change). This may indicate that *A. baumannii* adapts to cadmium stress by changing the structure and subsequently the biophysical properties of its membrane. Fatty acid analysis of *A. baumannii* under cadmium stress revealed minor but significant changes in 16:1 (1.5% decrease) and 18:0 (0.9% increase) (**Fig. S2**). These changes may indicate a slight increase in the membrane rigidity following cadmium exposure.

### *A. baumannii* employs a dedicated inner membrane cadmium exporter

To confirm the roles of the putative cadmium transporters in cadmium stress resistance as observed by TraDIS (**Fig. S3**), we examined mutants in *czcA* (ABUW_0268), *czcD* (ABUW_0269), *czcE* (ABUW_2851) and *czcF* (ABUW_3664). *A. baumannii* AB5075_UW and its mutant derivatives were grown with increasing concentrations of cadmium (1, 5, 10, 20 or 30 μM CdCl_2_) (**Fig. 2A-E**). We found that inactivation of *czcE* rendered the cells hyper-susceptible to cadmium stress, with significant growth perturbation seen at concentrations as low as 1 μM cadmium, which is approximately 30-fold lower than the AB5075_UW parental strain. In addition to a primary role in zinc resistance (22), CzcCBA contributes to cadmium resistance, with the *czcA*::T26 mutant showing increased sensitivity at 20 μM cadmium or greater. Consistent with the examination of these efflux systems in strain AB50875_UW, the ATCC 17978 Δ*czcA* and Δ*czcE* mutants displayed increased susceptibility to cadmium stress (**Fig. S4**). CzcD and CzcF were found not to play a major role in cadmium resistance. The concerted efforts from this study and previous work (22) have identified major roles for CzcA, CzcD and CzcE in zinc and/or cadmium resistance, but no major role has been identified for CzcF. Hence, we examined the response of CzcF to stress induced by other transition metals (zinc, copper, nickel, cobalt and iron), which revealed a minor role in zinc and nickel resistance (**Fig. S5**).

**Figure 2.**
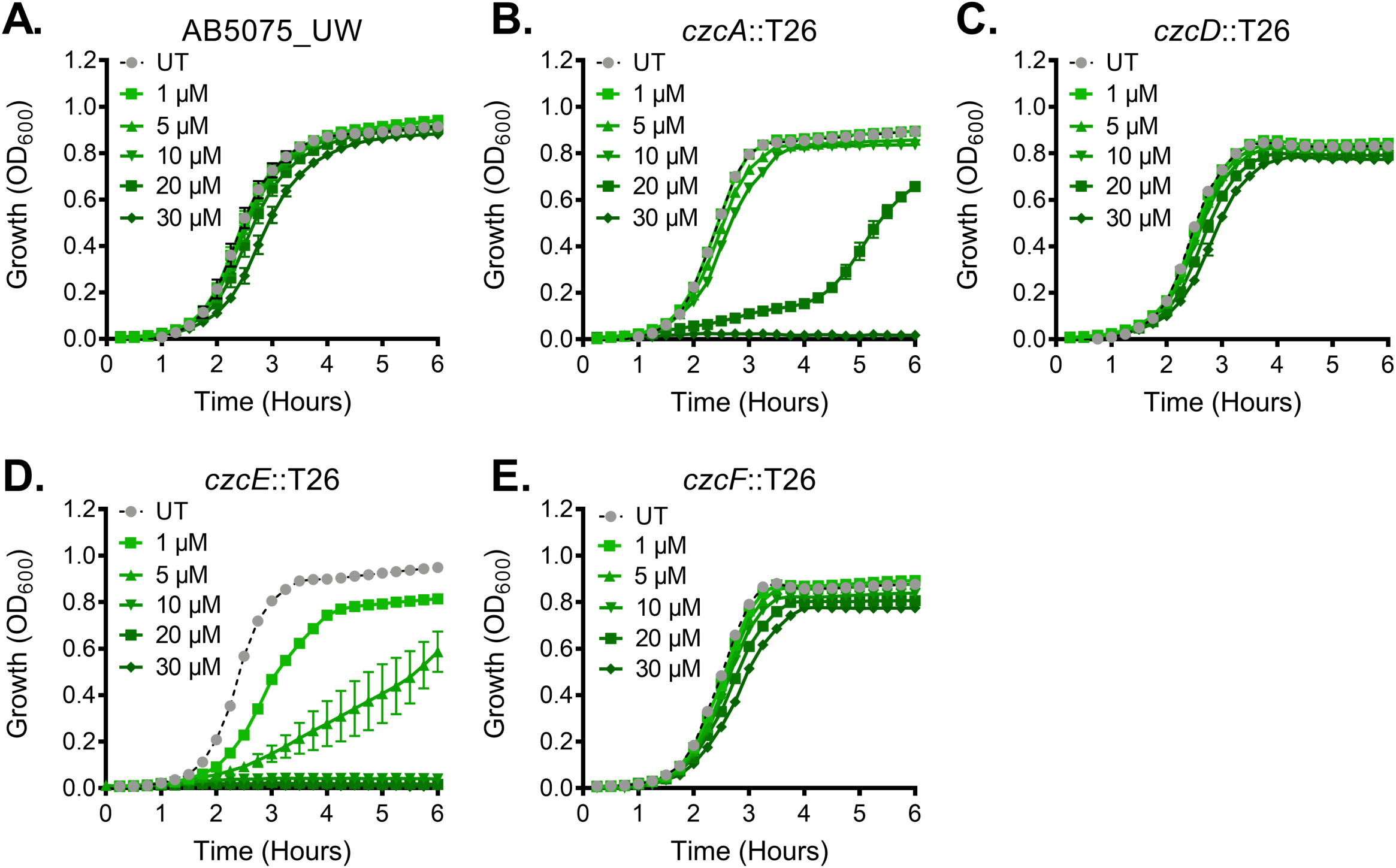
Cadmium resistance in *A. baumannii*. The effects of cadmium (1, 5, 10, 20 or 30 μM CdCl_2_) upon growth of the **(A)** wild-type, **(B)** *czcA*::T26, **(C)** *czcD*::T26, **(D)** *czcE*::T26 or **(E)** *czcF*::T26 mutant cells was compared to that of untreated (UT) cells. The growth was examined by measuring the optical density at 600 nm (OD_600_). The data are the mean of at least biological triplicates (± SEM). The visibility of error bars may be occluded by the symbols.

### The impact of cadmium on the *A. baumannii* metallome

Bacterial cells necessitate diligent balancing of metal ions as their hyper-accumulation or deficit beyond the cellular setpoints has detrimental impacts on metalloprotein function and has the potential to affect the homeostasis of other metals. First, we examined cadmium accumulation in *A. baumannii* strain AB5075_UW and the *czcA*::T26 and *czcE*::T26 mutant strains. These analyses revealed that CzcA inactivation did not impact accumulation following supplementation with 1 μM cadmium. In contrast, the *czcE*::T26 mutant accumulated cadmium at nearly 8-fold higher levels compared to the wild-type (**Fig. 3A**). Since growth of the *czcE*::T26 mutant was dramatically perturbed at concentrations greater than 1 μM cadmium, only the wild-type and *czcA*::T26 mutant were examined at 15 μM cadmium. The *czcA*::T26 mutant accumulated cadmium at greater levels than those observed for the wild-type (144 vs. 86 μg cadmium.g cells^-1^) (**Fig. 3A**).

**Figure 3.**
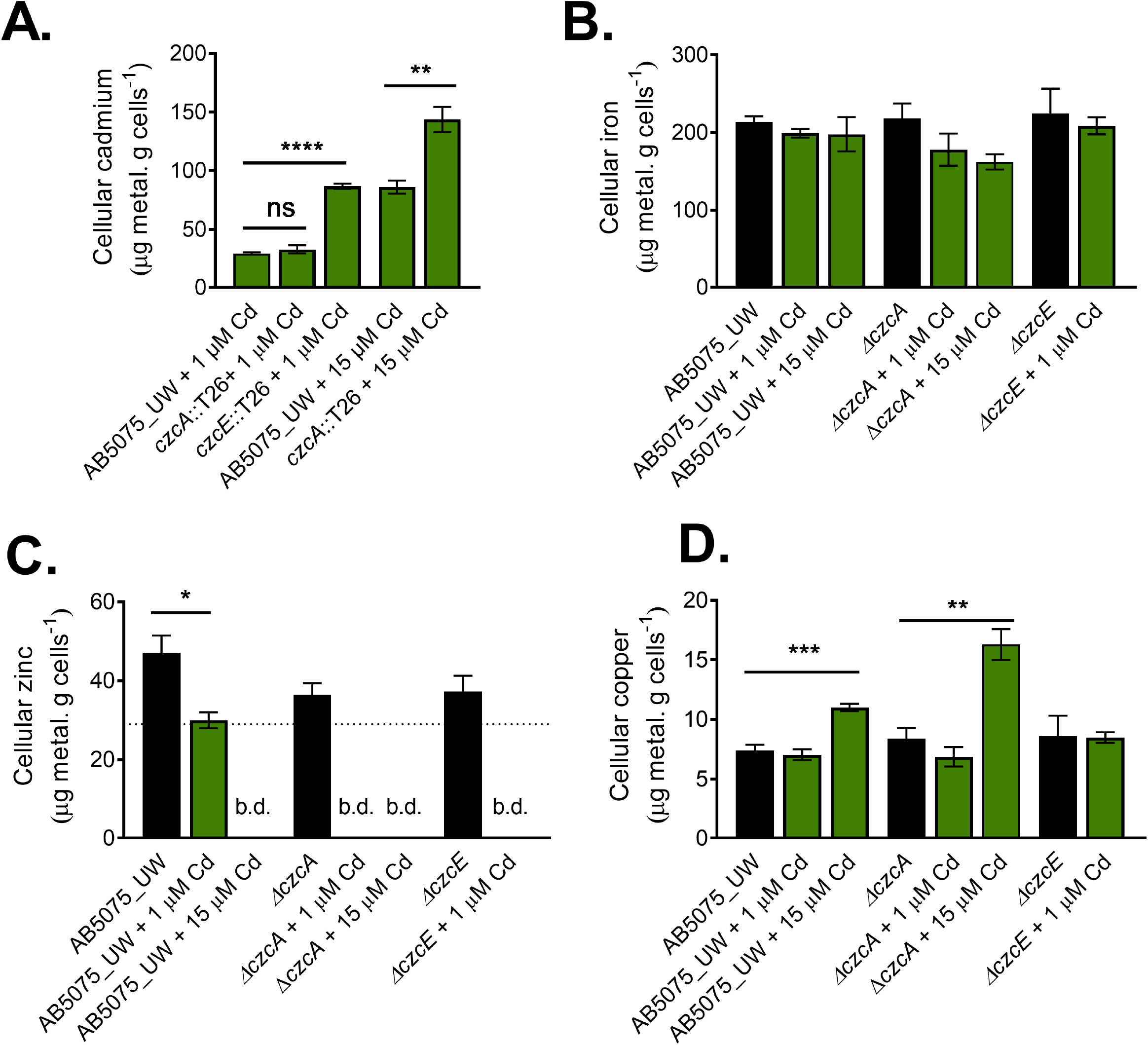
The impact of cadmium exposure on *A. baumannii* metal ion homeostasis. The accumulation of **(A)** cadmium, **(B)** iron, **(C)** zinc and **(D)** copper in wild-type, *czcA*::T26 or *czcE*::T26 cells with or without cadmium supplementation (1 μM or 15 μM CdCl_2_), as determined by inductively coupled plasma – mass spectrometry. Metal ion levels were quantified as the weight of the metal (μg) per the weight of the dry cell pellet (g). For all panels, the data are the mean of at least biological triplicates (± SEM). Statistical analyses were performed using a two-tailed Student’s *t*-test; n.s. = not significant, * = p < 0.05, ** = p < 0.01 and **** = p < 0.0001. The detection limit of zinc is indicated by the dotted line (b.d. = below detection).

Examination of the impact of cadmium on the most abundant *A. baumannii* metal ions revealed that zinc and copper levels were affected, but not iron (**Fig. 3B-D**). Zinc levels in the wild-type were significantly lower upon exposure to 1 μM cadmium and below detection when exposed to 15 μM cadmium. Consistent with these findings, zinc levels in the *czcA*::T26 and *czcA*::T26 mutants decreased to a greater extent upon exposure to 1 μM cadmium compared to the wild-type (**Fig. 3B**). Interestingly, the wild-type and *czcA*::T26 mutant hyper-accumulated copper when exposed to 15 μM cadmium (**Fig. 3D**). Despite the similar cellular cadmium levels observed in the *czcE*::T26 mutant following treatment with 1 μM cadmium and the wild-type with 15 μM cadmium (**Fig. 2A**), the accumulation of copper was not affected in the *czcE*::T26 mutant (**Fig. 2D**).

### Cadmium stress influences the *A. baumannii* metalloregulator responses

Transcriptional analyses revealed that *czcE* is up-regulated 480-fold by cadmium stress (**Fig. 4A**), while *czcI* was significantly down-regulated by cadmium stress. Analysis of CadR (ABUW_2852), a MerR-type regulator divergently transcribed from *czcE*, suggested that it was responsible for activation of *czcE*, which is consistent with the TraDIS analysis (**Fig. 4B**). We found that *czcF* was significantly up-regulated upon treatment with cadmium, despite it not playing a major role in cadmium resistance (**Fig. 4I and S6B**). Similar to *czcE*, we found that *czcF* is regulated by a divergently transcribed MerR-type regulator, ABUW_3665 (**Fig. 4C**).

**Figure 4.**
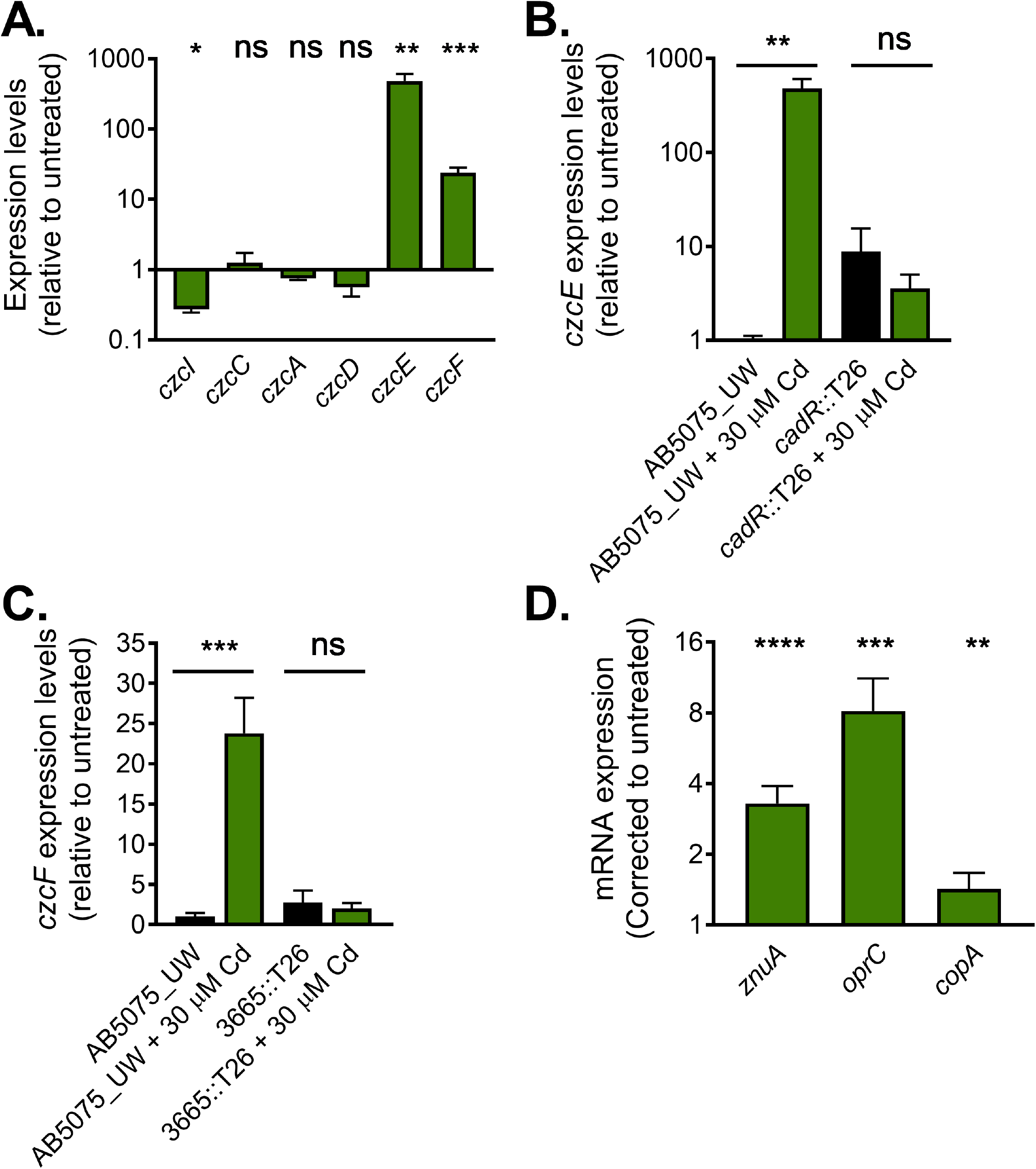
Regulation of cadmium resistance. **(A)** The transcriptional responses of *czcI, czcC, czcA, czcD, czcE* and *czcF* in wild-type AB5075_UW cells treated with 30 μM cadmium were examined by qRT-PCR. The transcription levels of **(B)** *czcE* or **(C)** *czcF* were examined in regulator mutant and AB5075_UW wild-type cells by qRT-PCR following treatment with 30 μM cadmium. **(D)** The transcription levels of *znuA, oprC* and *copA* in wild-type cells were examined by qRT-PCR following treatment with 30 μM cadmium. For all panels, the data are the mean (±SEM) of at least biological triplicates. The visibility of error bars may be occluded by the symbols. Statistical analyses were performed using a two-tailed Student *t*-test; n.s. = not significant, * = p < 0.05, ** = p < 0.01 and *** = p < 0.001.

Our metallomic analyses identified that cadmium toxicity impacts zinc and copper homeostasis (Fig. 2). To ascertain the impact of cadmium stress on the expression of metal ion homeostasis contributors, we analysed transcription of *znuA*, a component of the major zinc acquisition system ZnuABC (15), *oprC*, a putative copper acquisition system (43), and *copA*, the primary copper efflux system (22). Under cadmium stress, *A. baumannii* increases expression of *znuA* (3.3 log2-fold) (Fig. 4D), possibly in an attempt to compensate for zinc starvation. Interestingly, cadmium-induced copper overaccumulation may be a result of the significant up-regulation of *oprC* (8.2 log2-fold) (Fig. 4D).

## CONCLUSION

Cadmium resistance in *A. baumannii* is primarily mediated by a distinct CDF member, CzcE. However, mutation of *czcA* results in *A. baumannii* growth perturbation in the presence of cadmium, which suggests that cytoplasmic cadmium is translocated into the periplasm by CzcE and then exported into the extracellular space by CzcCBA. These findings on the roles of CDFs in conjunction with HME systems in the cytoplasmic and periplasmic translocation of substrates, respectively, are consistent with work performed with *Wautersia metallidurans* (44). Further, previous works have indicated that cadmium exerts greater toxicity in the cytoplasm compared to that of the periplasm (45, 46), which highlights the major significance of CzcE, and the lesser contribution of CzcA in *A. baumannii* cadmium resistance. Interestingly, transcriptional profiling revealed that *czcE* is transcriptionally activated by cadmium, whereas *czcA* transcription is not. Hence, the degree of CzcCBA contribution to cadmium resistance may also be due to the lack of up-regulation, rather than an inability of export from either the cytoplasm or periplasm.

Cadmium stress also results in copper over-accumulation with transcriptional analyses revealing *oprC* to be a possible copper import system. Although *oprC* has been shown to be regulated by OxyR in *Acinetobacter oleivorans* (47), recent analyses of OxyR in *A. baumannii* did not identify such association (48). Hence, the physiological role of OprC warrants further investigation. Zinc homeostasis was also affected by cadmium stress, resulting in a reduction in cellular zinc. However, the molecular basis behind cadmium-induced zinc depletion in *A. baumannii* remains unclear. These results show that cadmium stress results in an inverse dysregulation of copper and zinc homeostasis, which at is at least in part consistent with our previous analysis of these metals (13).

In summary, this study has identified the primary cadmium efflux pathways of *A. baumannii*. These systems are highly conserved across this species and genus, hence this work is broadly applicable to *Acinetobacter* biology, exemplified by our parallel analysis of two distinct strains. We have garnered new knowledge into the environmental stress adaptation strategies, with systems that have unique and shared metal ion specificities. The analyses of cadmium stress across a range of distinct metal ion export systems has also provided novel insights into the alignment of metal sensors and efflux systems, which is of major significance to bacterial fitness in a broad range of environments.

## Authors’ contributions

BAE, KAH, AKC, ITP, JCP and CAM designed the study. SFA, FGA, RM, KAH, NND and BAE performed the experiments. BAE, FGA, SFA, KAH, AKC, RM, ITP, CAM and JCP contributed to the drafting of the manuscript.

