## Supplemental Material for "The molecular basis of *Acinetobacter baumannii* cadmium toxicity and resistance"

**Table S1. Strains included in the study.**

| Name | Details | Origin |
|---|---|---|
| AB5075_UW | Wild-type | |
| *czcA*::T26 | tnab1_kr130913p05q135 | (1) |
| *czcD*::T26 | tnab1_kr130917p09q147 | (1) |
| *czcE*::T26 | tnab1_kr121128p05q140 | (1) |
| *czcF*::T26 | tnab1_kr121204p05q155 | (1) |
| *cadR*::T26 | tnab1_kr121128p05q189 | (1) |
| 3665::T26 | tnab1_kr121203p05q121 | (1) |
| ATCC 17978 | Wild-type | |
| ATCC 17978 + pAT04 | IPTG-inducible recombinase plasmid | (2) |
| 17978Δ*czcA* | Erythromycin$^{R}$ | (3) |
| 17978Δ*czcE* | Erythromycin$^{R}$ | (3) |

**Table S2. Oligonucleotides included in the study.**

| Name | Forward (5’ -> 3’) | Reverse (5’ -> 3’) |
|---|---|---|
| qABUW_0265 (czcI) | AATATAGCAGCGGCTTTTTGC | ATGGTCTTGTAAGTTTAAAGGT |
| qABUW_0266 (czcC) | AATGCAGTTGCAAGATGTGC | GTTGGAAAAGTTTGAGCTTG |
| qABUW_0268 (czcA) | GCTGCAACTTACAACGGTGA | TTTGTTTACTAAGCTCGTCC |
| qABUW_0269 (czcD) | TGCTTTGGTTGCCATACAGA | ACACTTTGAATTTCAGGTGG |
| qABUW_2851 (czcE) | AGGTGAAACGCCATGAAAAA | GCAGATGCATTAGATTTTGC |
| qABUW_3664 (czcF) | ATCCAGCTTGCCTTCATGTG | GCTGTGGGCGTACTCATAGA |
| qABUW_3740 (znuA) | GGGGCTGCGCTACCAAATAC | GCCGCCTGTAACATATTTCG |
| qABUW_3719 (oprC) | TCACCTGCCCTTCAATTTTC | GCTTGGCTTTACTCCAGATG |
| qABUW_2707 (copA) | TGGTTGCCGTTGATAAAACA | CTGCTTGAACAATCGCAAGG |
| mABUW_0268 (*czcA*::T26) | CTGCTTGAACAATCGCAAGG | CCTTTAACTCTTACAAACAC |
| mABUW_0269 (*czcD*::T26) | GGTGGACATCATGGTCATGATCATAG | ATGCTGATGTGAATGCGTCTTGTC |
| mABUW_2851-2 (*czcE*::T26 & *cadR*::T26) | GGAGTATTGCAATTTCACTTCTC | CCAGATGAACATAGCGATCG |
| mABUW_3664-5 (*czcF*::T26 & 3665::T26) | TACGGTGACAGATGTTTGCC | CGGCATCTCATCGGATTATC |
| CV_Tn5pUTKm | CTGTCTCTTATACACATCTGCCACGTTGTGTCTCAAAATCTC | CTGTCTCTTATACACATCTTCCCGTCAAGTCAGCGTAAGC |

| | | |
|---|---|---|
| Pf5_PCR | AATGATACGGCGACCACCGAGATCTACACATGATGATATATTTTTATCTTGTGCAATGTAACATC | AATGATACGGCGACCACCGAGATCTACACTCAGAATTGGTTAATTGGTTGTAACACTGGC |
| Pf5_5'Seq | CAGAGATTTTGAGACACAACGTGGCAGATGTGTA | GAGCATTACGCTGACTTGACGGGAAGATGTGTA |

24

**Table S3. TraDIS analyses of *A. baumannii* exposed to cadmium stress.**

| Locus tag | Function | Fold change ($Log_2$) | P value | Q value |
|---|---|---|---|---|
| ACX60_13165 | MerR family transcriptional regulator (CadR) | -2.05 | 8.6E-17 | 9.2E-15 |
| ACX60_13160 | CDF transport system (CzcE) | -1.94 | 9.0E-11 | 5.6E-09 |
| ACX60_09920 | biopolymer transporter ExbB | -1.94 | 8.7E-03 | 1.3E-01 |
| ACX60_10710 | methionine transporter | -1.66 | 1.9E-03 | 3.7E-02 |
| ACX60_15420 | lipoprotein | -1.61 | 1.4E-43 | 2.2E-40 |
| ACX60_01410 | hemolysin D | -1.60 | 1.8E-32 | 3.9E-30 |
| ACX60_01400 | cation transporter | -1.59 | 3.5E-34 | 8.4E-32 |
| ACX60_18405 | hypothetical protein | -1.54 | 4.2E-03 | 7.2E-02 |
| ACX60_01415 | cation transporter | -1.47 | 2.1E-34 | 5.4E-32 |
| ACX60_09235 | succinyl-CoA:3-ketoacid-CoA transferase | -1.31 | 1.9E-03 | 3.7E-02 |
| ACX60_01405 | RND transporter | -1.26 | 1.9E-16 | 1.8E-14 |
| ACX60_10905 | nitrate reductase | -1.23 | 2.3E-03 | 4.4E-02 |
| ACX60_18310 | IncC protein | -1.22 | 7.2E-02 | 4.7E-01 |
| ACX60_09155 | AraC family transcriptional regulator | -1.19 | 8.1E-05 | 2.5E-03 |
| ACX60_17750 | glucose-6-phosphate isomerase | -1.11 | 3.0E-21 | 4.5E-19 |
| ACX60_11665 | type VI secretion protein | -1.10 | 5.5E-02 | 4.1E-01 |
| ACX60_17830 | protein tyrosine phosphatase | -1.08 | 5.5E-06 | 2.0E-04 |
| ACX60_06050 | transporter | -1.07 | 2.6E-02 | 2.5E-01 |
| ACX60_07490 | hypothetical protein | -1.04 | 9.4E-02 | 5.3E-01 |
| ACX60_04945 | alanine acetyltransferase | -1.04 | 2.5E-02 | 2.5E-01 |
| ACX60_16900 | D-alanyl-D-alanine endopeptidase | -1.01 | 3.9E-17 | 4.3E-15 |
| ACX60_07585 | glutamate 5-kinase | -1.01 | 3.1E-02 | 2.9E-01 |
| ACX60_09300 | sulfonate ABC transporter ATP-binding protein | -1.00 | 7.9E-02 | 4.9E-01 |
| ACX60_16755 | DNA-binding protein | 3.66 | 1.2E-23 | 1.9E-21 |

| | | | | |
|---|---|---|---|---|
| ACX60_15250 | chemotaxis protein CheY | 3.60 | 9.2E-26 | 1.7E-23 |
| ACX60_16905 | response regulator | 3.55 | 6.1E-23 | 9.4E-21 |
| ACX60_05355 | peptidase S49 | 3.30 | 4.0E-35 | 1.2E-32 |
| ACX60_16845 | molecular chaperone DnaK | 3.09 | 1.2E-30 | 2.6E-28 |
| ACX60_09775 | membrane protein | 2.90 | 2.3E-28 | 4.4E-26 |
| ACX60_13270 | membrane protein | 2.85 | 4.3E-41 | 3.3E-38 |
| ACX60_06555 | mammalian cell entry protein | 2.82 | 6.2E-14 | 4.8E-12 |
| ACX60_01360 | fatty acyl-CoA reductase | 2.74 | 1.9E-39 | 9.8E-37 |
| ACX60_15855 | TetR family transcriptional regulator | 2.18 | 2.0E-40 | 1.3E-37 |
| ACX60_14885 | peptidase C13 family protein | 2.12 | 2.8E-39 | 1.2E-36 |
| ACX60_03910 | alcohol dehydrogenase | 2.09 | 8.3E-43 | 8.6E-40 |
| ACX60_03635 | acetyltransferase | 1.98 | 1.2E-37 | 4.5E-35 |
| ACX60_03915 | hypothetical protein | 1.93 | 3.7E-36 | 1.3E-33 |
| ACX60_01070 | monoamine oxidase | 1.85 | 5.3E-20 | 6.9E-18 |
| ACX60_03170 | folylpolyglutamate synthase | 1.80 | 4.0E-11 | 2.5E-09 |
| ACX60_05275 | TetR family transcriptional regulator | 1.74 | 3.5E-44 | 1.1E-40 |
| ACX60_00705 | hypothetical protein | 1.66 | 9.9E-35 | 2.8E-32 |
| ACX60_13350 | LysR family transcriptional regulator | 1.63 | 1.8E-18 | 2.2E-16 |
| ACX60_06745 | phosphoenolpyruvate synthase | 1.57 | 1.0E-14 | 8.2E-13 |
| ACX60_04580 | hypothetical protein | 1.56 | 5.0E-09 | 2.5E-07 |
| ACX60_15640 | peptidase S41 | 1.51 | 6.1E-21 | 8.5E-19 |
| ACX60_05075 | hypothetical protein | 1.51 | 3.2E-04 | 8.1E-03 |
| ACX60_06895 | 2-octaprenyl-3-methyl-6-methoxy-1,4-benzoquinol hydroxylase | 1.49 | 2.0E-18 | 2.4E-16 |
| ACX60_16435 | hypothetical protein | 1.47 | 3.7E-16 | 3.3E-14 |
| ACX60_03105 | acyl-CoA desaturase | 1.45 | 7.2E-21 | 9.6E-19 |
| ACX60_06330 | 30S ribosomal protein S21 | 1.42 | 1.2E-25 | 2.1E-23 |
| ACX60_00650 | DNA topoisomerase IV subunit B | 1.29 | 2.2E-08 | 1.1E-06 |
| ACX60_03825 | adenine deaminase | 1.28 | 1.0E-16 | 1.0E-14 |
| ACX60_05175 | isocitrate dehydrogenase | 1.23 | 3.2E-16 | 2.9E-14 |

| | | | | |
|---|---|---|---|---|
| ACX60_06715 | protoheme IX farnesyltransferase | 1.21 | 9.3E-03 | 1.3E-01 |
| ACX60_17425 | GMP synthase | 1.13 | 6.9E-03 | 1.1E-01 |
| ACX60_05695 | membrane protein | 1.12 | 9.1E-10 | 5.0E-08 |
| ACX60_16430 | hypothetical protein | 1.11 | 2.1E-11 | 1.4E-09 |
| ACX60_15950 | lauroyl acyltransferase | 1.10 | 5.9E-15 | 5.0E-13 |
| ACX60_18165 | cobalt transporter | 1.10 | 1.7E-09 | 8.8E-08 |
| ACX60_01140 | carboxymuconolactone decarboxylase family protein | 1.09 | 2.7E-07 | 1.1E-05 |
| ACX60_17860 | nicotinate-nucleotide pyrophosphorylase | 1.09 | 9.2E-09 | 4.5E-07 |
| ACX60_01930 | hypothetical protein | 1.07 | 3.3E-14 | 2.6E-12 |
| ACX60_13660 | IclR family transcriptional regulator | 1.07 | 3.7E-11 | 2.4E-09 |
| ACX60_16885 | thiol:disulfide interchange protein | 1.05 | 2.6E-18 | 3.0E-16 |
| ACX60_03705 | 3-methylitaconate isomerase | 1.05 | 7.7E-13 | 5.7E-11 |
| ACX60_17890 | TetR family transcriptional regulator | 1.02 | 1.3E-16 | 1.3E-14 |
| ACX60_00510 | nicotinate phosphoribosyltransferase | 1.00 | 2.2E-16 | 2.0E-14 |

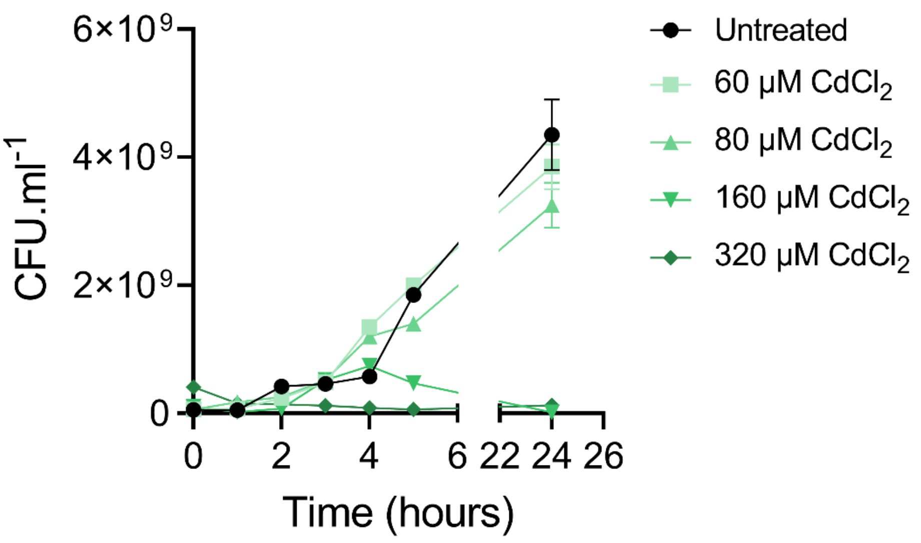

**Figure S1. Defining the optimal cadmium concentration for TraDIS analyses.** Overnight cultures of *A. baumannii* strain ATCC 17978 were diluted to $10^8$ CFU.mL$^{-1}$ and cultured with or without cadmium (60, 80, 160 or 320 µM $CdCl_2$). CFUs were enumerated at various time-points following serial dilution and plating. The data are a representative of replicate experiments.

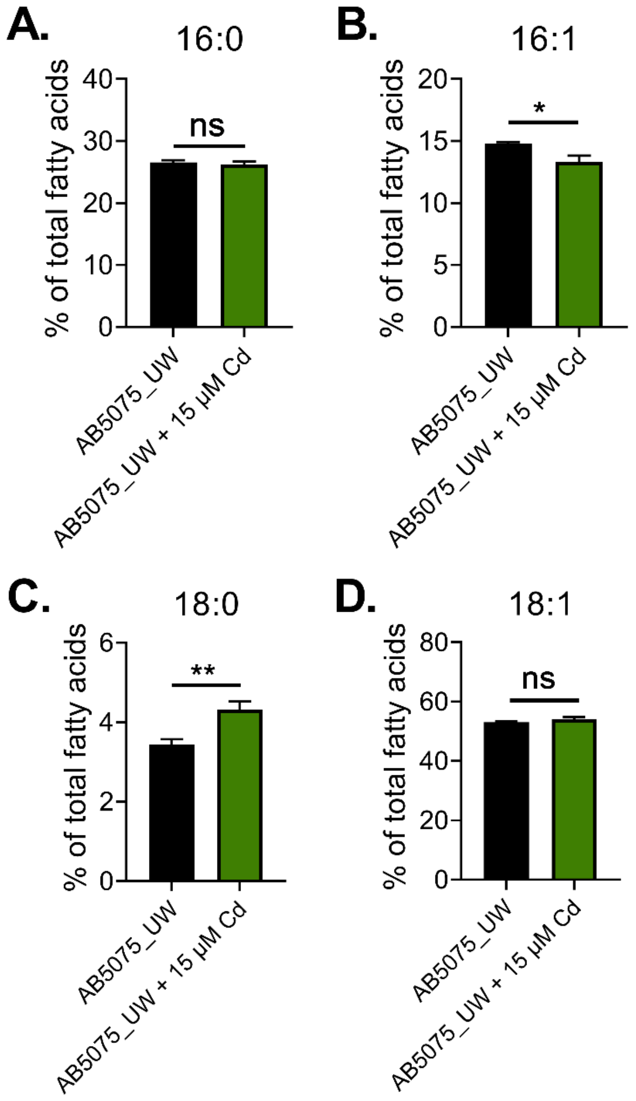

**Figure S2.** ***A. baumannii*** **fatty acid profiling in response to cadmium stress.** The *A. baumannii* AB5075_UW fatty acids **(A)** 16:0, **(B)** 16:1, **(C)** 18:0 and **(D)** 18:1 were quantified by gas chromatography/mass spectrometry following growth with and without supplementation with 15 µM $CdCl_2$. The data are the mean of at least biological triplicates (± SEM). Statistical analyses were performed using a two-tailed Student's *t*-test; n.s. = not significant, * = $p < 0.05$ and ** = $p < 0.01$.

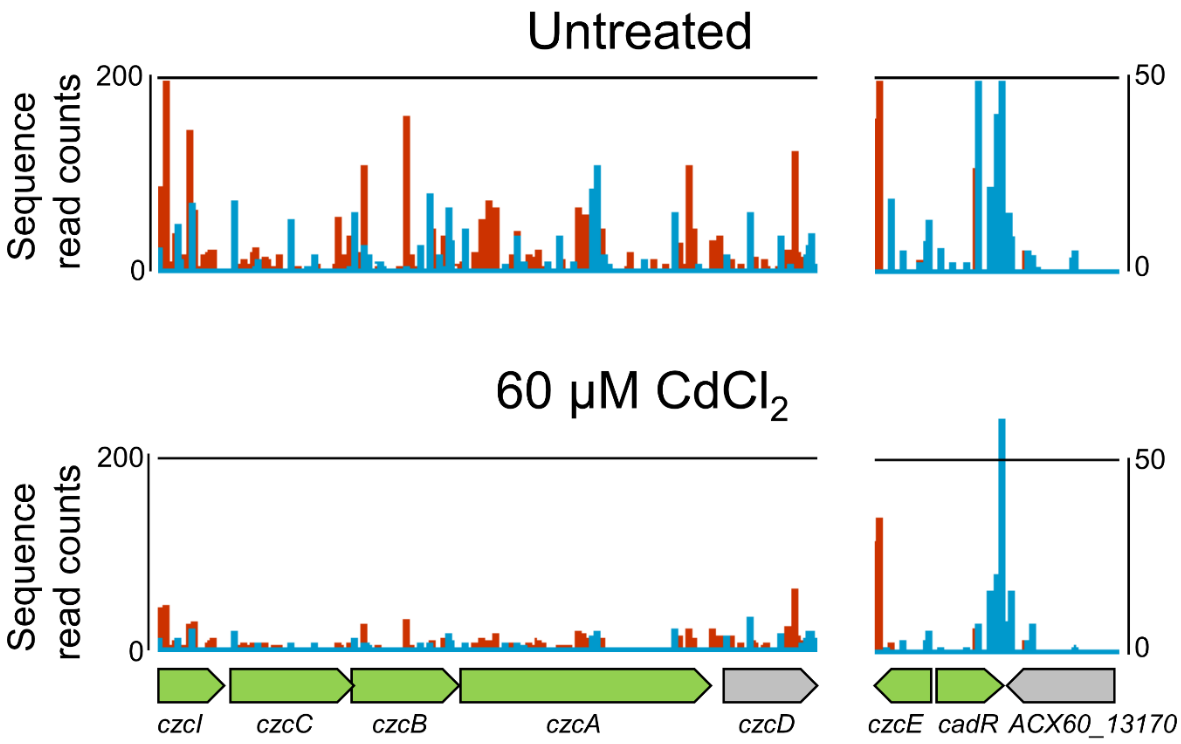

**Figure S3. TraDIS profiling of the *A. baumannii* metal ion transport systems.** Comparison of transposon (Tn*5*) insertion density between the untreated sample (MH control) and treatment with 60 µM $CdCl_2$ at the *czcICBAD* cation transport system and genes involved in MerR family transcriptional regulation, *czcE* and *cadR*. Vertical lines represent the density of transposon insertions at each insertion site, where red and blue indicate insertions in the forward or reverse direction, respectively. TraDIS analyses were performed in two independent cultures.

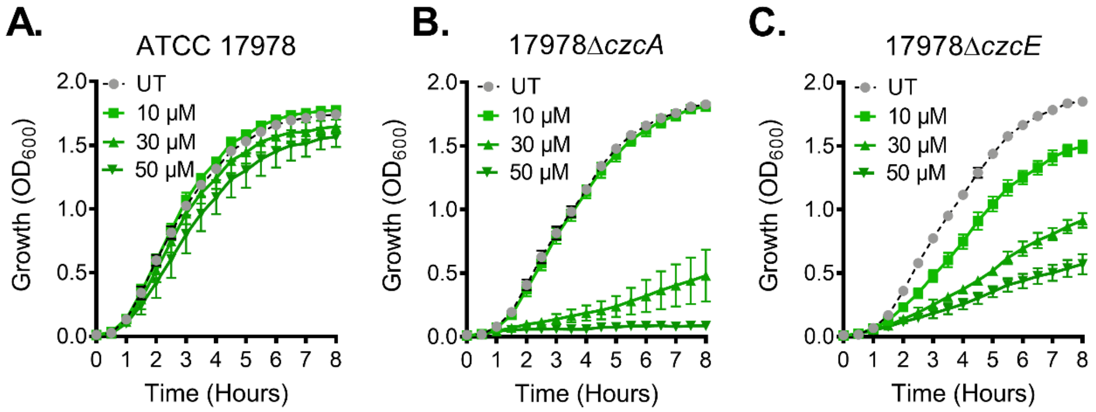

**Figure S4. Cadmium resistance in *A. baumannii* strain ATCC 17978.** The effects of cadmium (10, 30 or 50 µM) upon growth of the **(A)** ATCC 17978, **(B)** 17978Δ*czcA* or **(C)** 17978Δ*czcE* cells was determined by measuring the optical density at 600 nm ($OD_{600}$) every 15 min for 8 h. For all panels, the data represent the mean of at least biological triplicates (± SEM). The visibility of error bars may be occluded by the symbols. For all panels, the data are the mean of at least biological triplicates (± SEM).

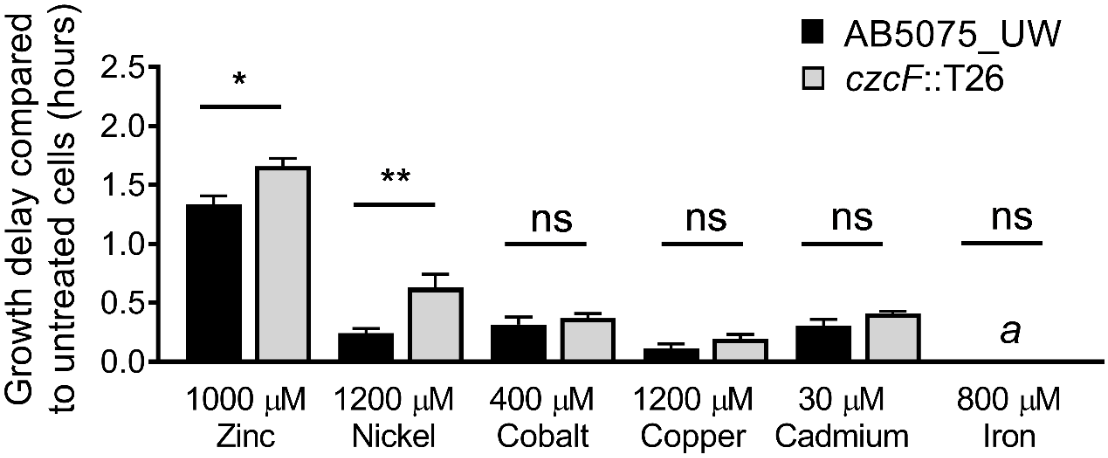

**Figure S5. Examination of CzcF under metal stress.** Growth of *A. baumannii* AB5075_UW and czcF::T26 mutant cells as determined by measuring the optical density at 600 nm (OD600). The growth delay was calculated following determination of the IC50 values, as described previously (4). Growth in the presence of 800 μM iron did not affect growth of the wild-type or *czcF*::T26 mutant (indicated with "*a*"), and was the highest concertation at which the metal remained soluble for at least 5 h. The data are the mean of at least biological triplicates (± SEM). Statistical analyses were performed using a two-tailed Student's *t*-test; n.s. = not significant, * = $p < 0.05$ and ** = $p < 0.01$.
